# c-MYC can Form Phase Condensates via its DNA Binding Domain

**DOI:** 10.1101/2025.04.22.650002

**Authors:** Udhaya Vijayakumar, Ruchi Choudhary, Wee-Joo Chng, Melissa Fullwood

## Abstract

The c-MYC oncogene is overexpressed in cancers. Phase condensates are membrane-less cellular compartments in cells. Using the optoDroplet system, which leverages optogenetics for condensate dynamics studies, here we showed c-MYC has the capability to form phase condensates. Moreover, the 3’ terminus of c-MYC, which contains a DNA binding domain but no intrinsic disordered regions, is sufficient to form phase condensates. Our research contributes to a better understanding of c-MYC biology.

## Introduction

The c-MYC transcription factor is crucial in numerous essential biological pathways, particularly in the abnormal growth and proliferation of neoplastic cells ^1,2^. It is noted that dysregulated c-MYC expression is observed in about 70% of human cancers, playing a significant role in their initiation and progression^3^. The regulation of c-MYC expression is complex, involving meticulous controls at transcriptional, post-transcriptional, translational, and post-translational levels. Regulatory enhancers and super-enhancers, which are often altered in cancerous cells, strictly control the transcription of the c-MYC proto-oncogene^4,5^. This widespread dysregulation of c-MYC, reflected in elevated mRNA and protein levels across various cancer types as documented in The Cancer Genome Atlas, underscores its potential as a target for therapeutic intervention^6^. However, the causes of c-MYC dysregulation are varied, ranging from gene amplification and mutations in oncogenic signalling pathways to structural genomic changes that introduce new regulatory elements near the MYC locus.

In normal cells, c-Myc serves as a critical link between growth factor stimulation and cellular proliferation^7^. Mitogenic signals increase MYC expression, with c-Myc boosting the transcription of genes associated with proliferation. However, in cancer cells where c-Myc is overexpressed, cellular proliferation becomes independent of growth factor signals, leading to the uncontrolled cell growth typical of cancer. This elevated c-Myc expression impacts various cellular functions, including chromatin structure, ribosome biogenesis, metabolic processes, cell adhesion, cell size, apoptosis, and angiogenesis. The mechanisms by which high levels of c-Myc lead to such diverse cellular changes remain unclear. Therefore, it becomes increasingly important to fully understand the various mechanisms by which c-MYC might regulate these numerous cellular processes.

Phase condensates, often referred to as biomolecular condensates, are cellular structures that form through a process called liquid-liquid phase separation (LLPS). These condensates are droplet-like compartments within the cell that lack a surrounding membrane, allowing molecules within them to concentrate and segregate from the bulk cellular environment^8^. This organization enables the cell to enhance biochemical reactions and regulate various cellular processes, including signal transduction, RNA processing, and stress response^9^. These are dynamic structures, which can reversibly form and dissolve depending on the cell’s needs and conditions. These condensates are composed of proteins and nucleic acids that exhibit multivalent interactions, driving the separation from the surrounding cytoplasm or nucleoplasm.

Recent studies have identified c-MYC multimeric spherical bodies that potentially shield replication forks^10^. Furthermore, recombinant c-MYC proteins have been shown to form droplets *in vitro* under specific conditions, such as the presence of LLPS buffers and appropriate salt concentrations^10,11^. However, these observations were largely based on *in vitro* droplet assays that may not mimic the actual cellular conditions. To understand the dynamics of phase separation in living cells, optoDroplet systems can be used. This method employs optogenetics to create and manipulate condensates in a highly controlled manner using light. OptoDroplets are generated by fusing proteins of interest with light-responsive domains that undergo oligomerization upon exposure to specific wavelengths of light. This light-induced oligomerization mimics the multivalent interactions that drive phase separation in biological systems. By exposing these fusion proteins to light^12^. This method can be used to rapidly induce the formation of condensates and subsequently dissolve them by turning off the light or using a different light wavelength, offering a reversible and highly tuneable system. Hence, in this manuscript, we aimed to understand the ability of c-MYC to form phase condensates using optoDroplet system and to identify the specific domains required for the formation in live cells.

## Results

### An optoDroplet system demonstrates that c-MYC can form phase condensates

For this study, we hypothesized that c-MYC can form dynamic phase condensates. To test this hypothesis, we employed the optoDroplet system. The optoDroplet system provides a valuable method for testing whether a specific protein can form condensates by genetically fusing the protein of interest to a light-responsive domain that undergoes oligomerization upon exposure to specific wavelengths of light. This fusion protein is then expressed in living cells. Upon light exposure, the light-responsive domain oligomerizes, mimicking the multivalent interactions necessary for phase separation. If the protein of interest can form condensates, the formation of optoDroplets can be observed using fluorescence microscopy.

We fused the light-sensitive protein Cry2, known for its ability to self-associate upon exposure to blue light, to the c-MYC protein (Fig.1a, b). U2OS cells expressing opto-cMYC (1-439) were exposed to blue light (488 nm pulsed laser) for 1 minute, and mCherry reporter signals were simultaneously captured using a Zeiss LSM 710 confocal microscope (Fig. 1c). Upon blue light exposure, we observed rapid clustering of human c-MYC in the nucleus (Fig. 1c, f, Supplementary Movie 1c). As a positive control, Opto-FUSN also exhibited optoDroplet formation, while Cry2-WT alone did not form droplets under identical exposure conditions (Fig. 1c, d, e, Supplementary Movie 1a, b). Furthermore, U2OS cells expressing Cry2-WT, Opto-FUSN, or opto-cMYC (1-439) and exposed to continuous blue light for 2 minutes under the Zeiss Live Cell Observer II exhibited similar results (Supplementary Fig. 1a). These findings suggest that c-MYC possesses an intrinsic propensity for self-assembly in the cellular environment. Upon withdrawal of blue light, opto-cMYC (1-439) clusters reverted to a diffuse state within 5 minutes (Fig. 1f, Supplementary Movie 1c) similar to that of positive control Opto-FUSN (Fig. 1e, Supplementary Movie 1b). This ability of c-MYC droplets to rapidly dissolve upon withdrawal of blue light indicates that c-MYC condensates are dynamic structures.

**Figure 1:**
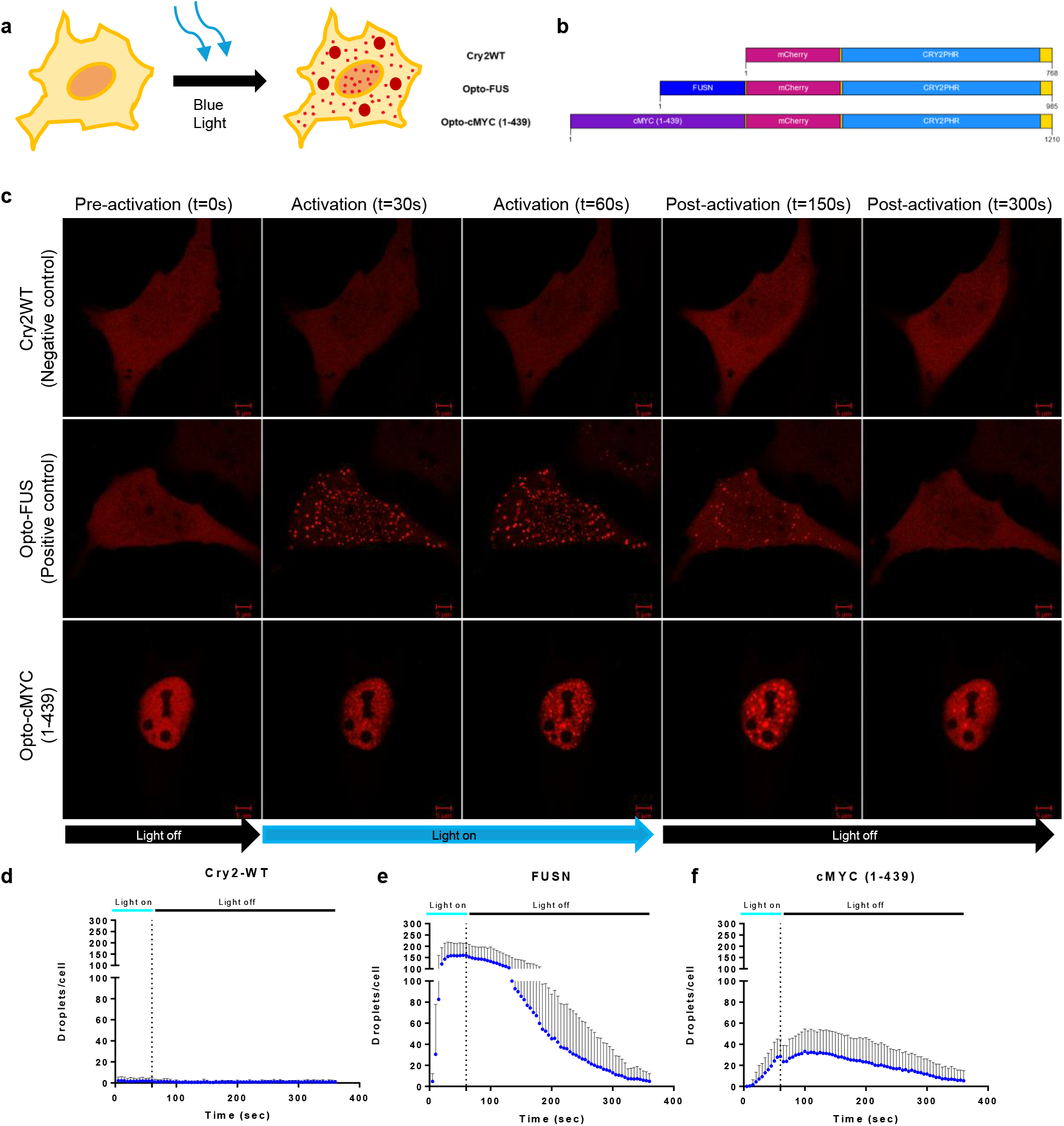
c-MYC undergoes phase separation upon blue light activation in the optoDroplet system. (a) Schematic diagram of the optoDroplet system, illustrating how blue light induces the clustering of target protein domains in cells. (b) Schematic illustration^12^ of the optoDroplet constructs, where opto-cMYC(1-439) consists of Cry2PHR, mCherry, and the full-length human cMYC. (c) Representative images of U2OS cells expressing mCherry-Cry2 alone or mCherry-Cry2 fused to FUSN or MYC(1-439), during blue light activation and deactivation. Cells were imaged at 5-second intervals under identical blue light activation conditions. Scale bar = 5 µm. (d-f) Quantification of the total number of droplets during blue light activation and deactivation for (d) Cry2WT, (e) opto-FUSN, and (f) opto-cMYC(1-439). Data are represented as mean ± SD, with n = 11 cells for Cry2WT, n = 12 cells for opto-FUSN, and n = 14 cells for opto-cMYC(1-439).

### The DNA Binding Domain of c-MYC is essential for phase condensate formation

Next, to identify which domain of the c-MYC protein is essential for phase condensate formation we fused Cry2 with four different c-MYC constructs: The first construct, opto-cMYC (1-169), covers the N-terminal transcriptional activation domain (TAD), while the second, opto-cMYC (204-327), includes the central region, predicted to be an intrinsically disordered region (IDR) based on iUPred Score^13^ (Fig. 2a, b). The third construct, opto-cMYC (334-439), covers the C-terminal DNA binding domain (Fig. 2a, b). All the constructs were sequence verified, and the protein expression length were verified by western blot (Supplementary Fig. 2) Results from optoDroplet assays revealed that only opto-cMYC (334-439) formed protein clusters, indicating self-interaction within the nucleus (Fig. 2c, f, Supplementary Movie 1f). In contrast, the opto-cMYC (1-169) and opto-cMYC (204-327) constructs did not form clusters in the cells (Fig. 2c, d, e, Supplementary Movie 1d, e) within 1 minute of pulsated blue light exposure. Additionally, after 2 minutes of continuous blue light exposure, the opto-cMYC (1-169) construct formed droplets, while opto-cMYC (204-327) still did not form clusters under Zeiss Live Cell Observer II (Supplementary Fig. 1b). Taken together, these findings suggest that the C-terminal DNA-binding domain is sufficient for c-MYC phase condensate formation.

**Figure 2.**
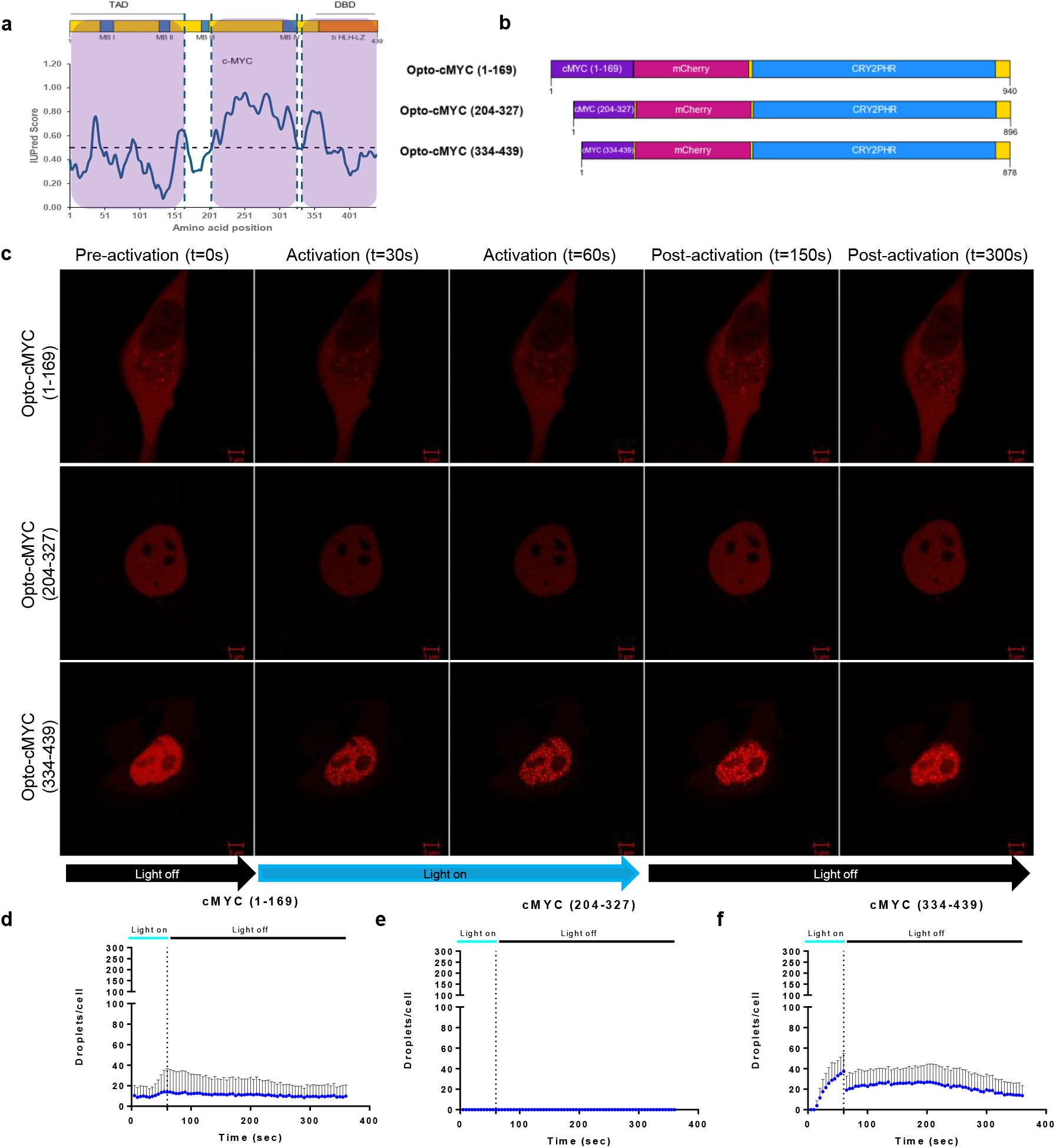
The DNA Binding Domain (DBD) of c-MYC undergoes phase separation upon blue light activation. (a) Domain structure^12^ of the c-MYC protein, with the Transactivation Domain (TAD), DNA Binding Domain (DBD), MYC boxes (MB), and basic helix-loop-helix leucine zipper (bHLHLZ) regions shown at the top. The blue line in the graph represents the IUPred score^11^ for intrinsically disordered regions of c-MYC, with values above 0.5 (indicated by the dotted black lines) considered disordered. Highlighted violet box are regions of c-MYC selected for opotdroplet study (b) Schematic illustration^12^ of the optoDroplet constructs. opto-cMYC(1-169) consists of Cry2PHR, mCherry, and the 1-169 amino acids (aa) of human cMYC. opto-cMYC(204-327) consists of Cry2PHR, mCherry, and the 204-327 aa of cMYC. opto-cMYC(334-439) consists of Cry2PHR, mCherry, and the 334-439 aa of cMYC. (c) Representative images of U2OS cells expressing mCherry-Cry2 fused to MYC(1-169), MYC(204-327), or MYC(334-439) during blue light activation and deactivation. All cells were activated under identical blue light conditions and imaged at 5-second intervals. Scale bar = 5 µm. (d-f) Quantification of the total number of droplets during blue light activation and deactivation for (d) opto-cMYC(1-169), (e) opto-cMYC(204-327), and (f) opto-cMYC(334-439). Data are presented as mean ± SD, with n = 11 cells for opto-cMYC(1-169), n = 12 cells for opto-cMYC(204-327), and n = 12 cells for opto-cMYC(334-439).

## Discussion

Our experimental results show that c-MYC can form phase condensates. This capability suggests that c-MYC might orchestrate various cellular transformations through its participation in these phase-separated structures. Our results are consistent with the observations reported in a recent study about c-MYC multimeric spherical bodies that shield the replication fork^10^. Such condensates and multimeric bodies likely create a concentrated and localized setting for interactions, crucial for numerous biological processes. This insight into c-MYC’s condensate formation unveils new possibilities for therapeutic strategies. Targeting the traditionally “undruggable” c-MYC via its phase condensates presents an innovative approach, circumventing the usual challenges of directly inhibiting this protein. This method could lead to novel treatments that disrupt the pathological condensates specifically, offering a focused way to tackle cancer cells driven by c-MYC dysregulation.

Second, we concluded that the C-terminal region of c-MYC, which contains a DNA binding domain (C-terminal) of c-MYC, but no IDRs, can form phase condensates. While many previous studies^14-16^ have emphasized the significant role of intrinsically disordered regions (IDRs) in phase condensate formation, our findings indicate that IDRs are neither necessary nor sufficient for the formation of condensates in the case of c-MYC. This observation aligns with findings related to other proteins, such as CTCF, where the DNA binding region also plays a crucial role in condensate formation^17^, similar to what we have observed with c-MYC.

Taken together, our findings not only enhance our understanding of c-MYC biology but also underscore the significance of the DNA binding regions in proteins for the formation of phase condensates.

## Methods

### Cell Culture

U2OS osteosarcoma cells (ATCC HTB-96), which contain a doxycycline-inducible MYC system, were kindly provided by Elmar Wolf’s lab at the Universität Würzburg, Germany. The cells were cultured in Dulbecco’s modified Eagle’s medium (DMEM, GIBCO) supplemented with 10% tetracycline-free fetal bovine serum (Clontech) and 100 U/mL penicillin-streptomycin (GIBCO). Cells were incubated at 37 °C in a humidified incubator with 5% CO2. The cells were maintained and sub-cultured every 2–4 days according to ATCC recommendations.

### Plasmid Construction

DNA fragments encoding human c-MYC (residues 1–439), human c-MYC (residues 1–169), human c-MYC (residues 204–327), and human c-MYC (residues 334–439) were amplified by polymerase chain reaction (PCR) using the plasmid pCMV4a-Flag-c-Myc (Addgene 102625) as the template. The pHR-mCh-Cry2WT plasmid (Addgene 101221) was linearized by PCR. The c-MYC DNA fragments were inserted into the linearized pHR-mCh-Cry2WT backbone using Gibson Assembly® Master Mix (New England Biolabs). The resulting constructs were Sanger sequenced to ensure the absence of unwanted mutations with the help of primers 5’ CCAGAGGTTGATTATCGATAAGC 3’ and 5’ GCTTCCCGAGCTCTATAAAAGAG 3’. Plasmids were then transformed into One Shot™ Stbl3™ Chemically Competent E. coli (Invitrogen) for amplification and storage.

### Transient Transfections

U2OS cells were plated and grown to 60% confluency prior to transfection. Cells were transfected with 250 ng of plasmid encoding the CRY2-mCherry constructs, along with 750 ng of polyethyleneimine (PEI) at a DNA to PEI ratio of 1:3. Cells were imaged 48 hours post-transfection.

### optoDroplet Imaging

Confocal images were acquired using a Zeiss LSM 710 confocal microscope equipped with a 63X oil immersion objective (NA 1.4). An imaging chamber was maintained at 37 °C and 5% CO2. For live-cell imaging, cells were seeded in µ-Slide 8 Well high Glass Bottom dishes (ibidi). For droplet formation, cells were activated with a 488-nm laser (0.1% power) and imaged using a 560-nm laser every 5 seconds for 1 minute. After 1 minute of blue light exposure, cells were imaged using the 560-nm laser every 5 seconds in the absence of blue light until 5 minutes.

### Puncta Image Analysis

To quantify optoDroplets of mCherry signals in light-activated condensates, images were analyzed using a custom-built CellProfiler pipeline. The primary objects representing the nucleus and cell were identified, and droplets were detected using enhanced or suppressed features and labeled as secondary objects. The intensity and count of the identified objects were measured and plotted.

### Statistics and Reproducibility

Results are presented as the mean ± standard deviation. All micrographs in this study are representative of experiments conducted with at least three biological replicates. Graphs were plotted using GraphPad Prism version 10.0.0 for Windows, GraphPad Software, Boston, Massachusetts USA, www.graphpad.com.

### Plot of optoDroplet construct, c-MYC domain and disorder score

The illustration of the optoDroplet construct and the c-MYC domain was generated using Illustrator for Biological Sequences 2.0 (IBS 2.0)^18^. To assess the intrinsic disorder of the c-MYC protein, amino acid disorder scores were calculated using IUPred3^11^.

### Sample preparation and Western Blot

Cell lysates were prepared by resuspending samples in RIPA buffer with protease inhibitor cocktail. After incubation on ice for 30 minutes, the lysates were cleared by centrifugation at 16,000 × g for 20 minutes at 4°C. The protein concentration was determined using the BCA protein assay kit (Pierce, Thermo Fisher Scientific) according to the manufacturer’s protocol, and the samples were adjusted to an equal concentration for subsequent analysis.

Protein samples (15 µg) were mixed with a 4× Laemmli sample buffer (Bio-Rad) and heated at 95°C for 5 minutes to denature the proteins. The samples were then resolved on 8% SDS-polyacrylamide gel and transferred onto an Immobilon-P membrane (Millipore). The membrane was blocked with 5% non-fat milk in Tris-buffered saline containing 0.1% Tween (TBST) for 1 h at room temperature, and then probed with mCherry polyclonal antibody (PA5-34974) (1:3000 dilution) overnight at 4 °C. After washing with TBST, the membrane was incubated with Anti-rabbit IgG, HRP-linked (7074P2, Cell signalling) (1:3000 dilution) for 1 h at room temperature. Washed blots were developed using Clarity Western ECL substrate (Biorad).

## Data availability

All the raw images can be found at DR-NTU(data) repository https://doi.org/10.21979/N9/DUJJ55

## Acknowledgements

This research is supported by the National Research Foundation Singapore under its Open Fund - Individual Research Grant (MOH-001387) and administered by the Singapore Ministry of Health’s National Medical Research Council, and by the Ministry of Education, Singapore, under its Academic Research Fund Tier 1 (RG38/23), both awarded to M.J.F. (PI).

## Author contributions

U.V., R.C and M.J.F conceived the research idea. U.V. designed the constructs and performed the experiments. R.C. advised on the data analyses for the experiments. R.C., U.V. and M.J.F wrote the manuscript. W.J.C and M.J.F supervised the research. All authors reviewed and approved the manuscript.

## References

1 Evan, G. I. & Littlewood, T. D. The role of c-myc in cell growth. Curr Opin Genet Dev 3, 44–49 (1993). 10.1016/s0959-437x(05)80339-9

2 Dang, C. V. c-Myc target genes involved in cell growth, apoptosis, and metabolism. Mol Cell Biol 19, 1–11 (1999). 10.1128/MCB.19.1.1

3 Arango, D. et al. c-Myc overexpression sensitises colon cancer cells to camptothecin-induced apoptosis. Br J Cancer 89, 1757–1765 (2003). 10.1038/sj.bjc.6601338

4 See, Y. X., Chen, K. & Fullwood, M. J. MYC overexpression leads to increased chromatin interactions at super-enhancers and MYC binding sites. Genome Res 32, 629–642 (2022). 10.1101/gr.276313.121

5 Affer, M. et al. Promiscuous MYC locus rearrangements hijack enhancers but mostly super-enhancers to dysregulate MYC expression in multiple myeloma. Leukemia 28, 1725–1735 (2014). 10.1038/leu.2014.70

6 Rau, A., Flister, M., Rui, H. & Auer, P. L. Exploring drivers of gene expression in the Cancer Genome Atlas. Bioinformatics 35, 62–68 (2019). 10.1093/bioinformatics/bty551

7 Greenberg, M. E., Hermanowski, A. L. & Ziff, E. B. Effect of protein synthesis inhibitors on growth factor activation of c-fos, c-myc, and actin gene transcription. Mol Cell Biol 6, 1050–1057 (1986). 10.1128/mcb.6.4.1050-1057.1986

8 Feng, Z., Chen, X., Wu, X. & Zhang, M. Formation of biological condensates via phase separation: Characteristics, analytical methods, and physiological implications. J Biol Chem 294, 14823–14835 (2019). 10.1074/jbc.REV119.007895

9 Li, X. H., Chavali, P. L., Pancsa, R., Chavali, S. & Babu, M. M. Function and Regulation of Phase-Separated Biological Condensates. Biochemistry 57, 2452–2461 (2018). 10.1021/acs.biochem.7b01228

10 Solvie, D. et al. MYC multimers shield stalled replication forks from RNA polymerase. Nature 612, 148–155 (2022). 10.1038/s41586-022-05469-4

11 Boija, A. et al. Transcription Factors Activate Genes through the Phase-Separation Capacity of Their Activation Domains. Cell 175, 1842–1855 e1816 (2018). 10.1016/j.cell.2018.10.042

12 Shin, Y. et al. Spatiotemporal Control of Intracellular Phase Transitions Using Light-Activated optoDroplets. Cell 168, 159–171 e114 (2017). 10.1016/j.cell.2016.11.054

13 Erdos, G., Pajkos, M. & Dosztanyi, Z. IUPred3: prediction of protein disorder enhanced with unambiguous experimental annotation and visualization of evolutionary conservation. Nucleic Acids Res 49, W297–W303 (2021). 10.1093/nar/gkab408

14 Shin, Y. & Brangwynne, C. P. Liquid phase condensation in cell physiology and disease. Science 357 (2017). 10.1126/science.aaf4382

15 Jo, Y., Jang, J., Song, D., Park, H. & Jung, Y. Determinants for intrinsically disordered protein recruitment into phase-separated protein condensates. Chem Sci 13, 522–530 (2022). 10.1039/d1sc05672g

16 Wang, Y. et al. Phase separation of SPIN1 through its IDR facilitates histone methylation readout and tumorigenesis. J Mol Cell Biol 16 (2024). 10.1093/jmcb/mjae024

17 Zhou, R. et al. CTCF DNA-binding domain undergoes dynamic and selective protein-protein interactions. iScience 25, 105011 (2022). 10.1016/j.isci.2022.105011

18 Xie, Y. et al. IBS 2.0: an upgraded illustrator for the visualization of biological sequences. Nucleic Acids Res 50, W420–W426 (2022). 10.1093/nar/gkac373

